# Alternative polyadenylation regulates human urothelial cell differentiation

**DOI:** 10.1101/2025.03.03.641311

**Authors:** Ninh B. Le, Surbhi Sona, Briana Santo, R. Allen Schweickart, Veena Kochat, William I. Padron, Kunal Rai, Shih-Han Lee, Joo Mi Yi, Oliver Wessely, Byron H. Lee, Angela H. Ting

## Abstract

The urothelium is stratified into progenitor basal cells, intermediate cells, and terminally differentiated umbrella cells. Proper renewal of umbrella cells is necessary for maintaining urinary tract barrier integrity. To investigate whether mRNA alternative cleavage and polyadenylation (APA) regulates urothelial differentiation, we developed a single-cell polyadenylation site usage (scPASU) computational pipeline to map cell state-specific polyadenylation sites in single-cell RNA-seq data from 13,544 urothelial cells. Leveraging single-cell spatial imaging, we directly visualized APA events *in situ*, revealing their spatial specificity within the adult human ureter. APA shaped urothelial differentiation, independent of gene expression changes. Furthermore, key APA-regulated genes shared conserved motifs in their 3’ UTRs, often containing *Alu* elements, suggesting a potential mechanism regulating poly(A) site selection. Our study establishes APA as a driver of urothelial transcriptome diversity.

---

The urothelium of the adult human ureter consists of the entire spectrum of epithelial differentiation from the progenitor basal cells to the terminally differentiated umbrella cells lining the lumen. Proper renewal of umbrella cells after injury is essential for maintaining the integrity of the ureter to provide passage of urine from the kidneys to the bladder. Cellular differentiation is driven by precise transcriptional and post-transcriptional regulation to ensure that the correct genes are expressed in the right place at the right time. Understanding the transcriptomic dynamics across the urothelial differentiation spectrum has broad implications for both tissue repair and disease pathogenesis. While a prior single cell RNA-seq (scRNA-seq) study revealed highly similar gene expression patterns among basal, intermediate, and umbrella cells, the role of other mechanisms that fine-tune the transcriptome has not been investigated ^1^. mRNA alternative cleavage and polyadenylation (APA) is a process where distinct mRNA 3’ ends are generated to provide transcript diversity that can influence message stability, localization, and translation under a variety of physiologic and pathophysiologic conditions. Here, we developed a computational pipeline to examine the role of APA in gene expression regulation during urothelial homeostasis. We found that APA is an important mechanism for regulating gene expression in urothelial differentiation. We also discovered that a subset of the differentiation associated APA genes share simian-conserved sequence motifs in their 3’ untranslated regions (UTRs) that might suggest primate specific regulation of these genes.

## 3’ scRNA-seq was repurposed for mapping and quantifying mRNA polyadenylation sites

To investigate whether APA shapes gene expression patterns during urothelial differentiation, we re-analyzed scRNA-seq data for 13,544 urothelial cells from 10 healthy human ureters^1^ to map and quantify urothelial cell-specific mRNA polyadenylation [poly(A)] sites. The 3’ scRNA-seq library, generated using the 10x Genomics protocol (**Fig. 1A**, left), was virtually identical to those produced by bulk poly(A) sequencing strategies (**Fig. 1A**, right). Both used anchored poly(dT) primers to capture the poly(A) tails of mRNA for cDNA synthesis, and the resulting sequencing reads were near poly(A) sites. There was, however, a key difference: conventional poly(A)-seq generated sequencing reads spanning the poly(A) junction, whereas 3’ scRNA-seq yielded R2 reads upstream of the actual poly(A) junction by an offset equivalent to the average library fragment size. Thus, we developed a computational pipeline for single cell poly(A) site usage (scPASU), based on a previous bulk poly(A)-seq analysis workflow^2^, while accounting for the positional offset (**Fig. 1B**).

**Fig. 1.**
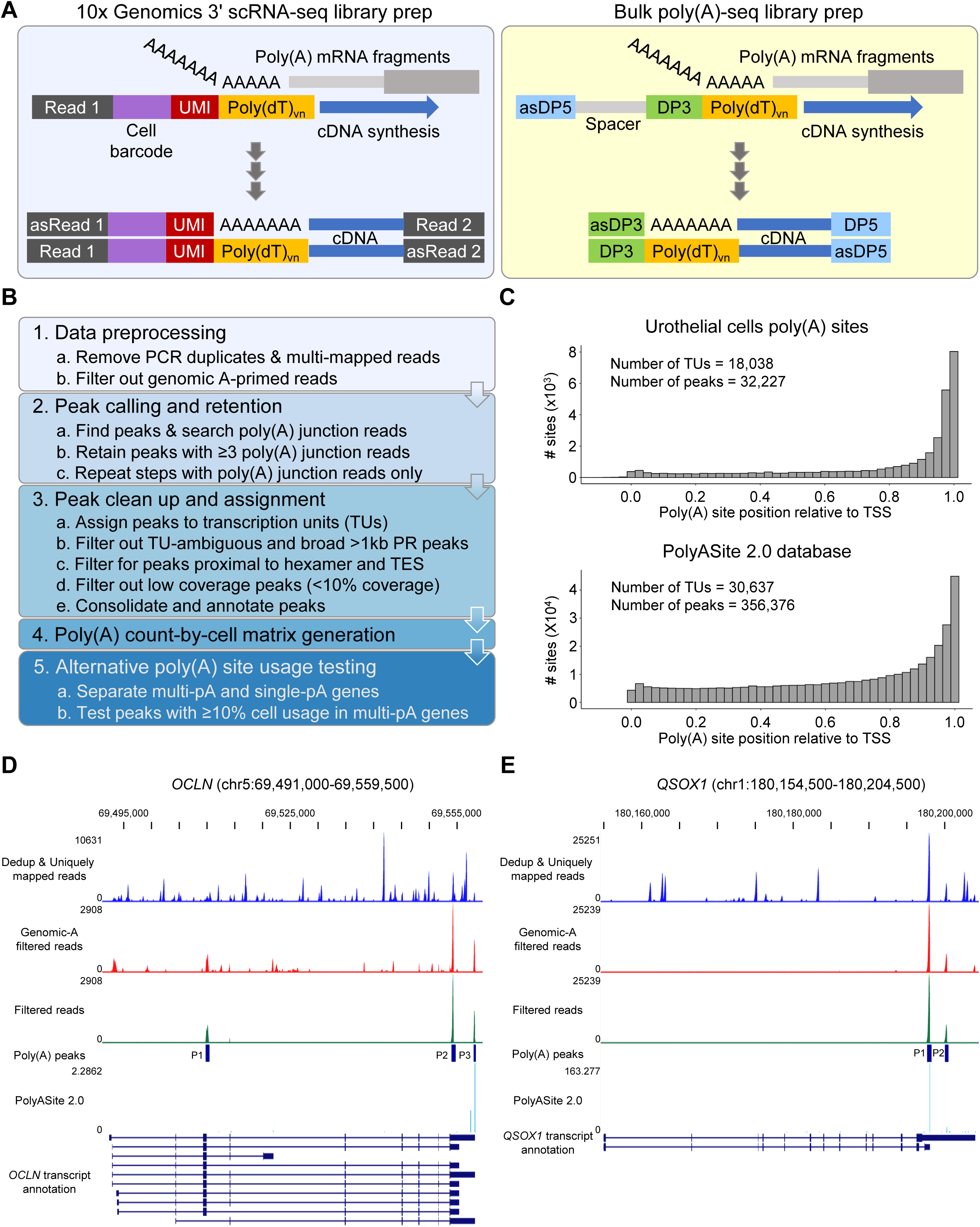
Urothelial-specific poly(A) site usage can be extracted from standard 3’ scRNA-seq data. (**A**) Schematic showing the sequencing library preparations for 10x Genomics 3’ scRNA-seq (left) and bulk poly(A)-seq (right). UMI is the unique molecular identifier used for identifying PCR duplicates. DP3 and DP5 are Illumina sequencing adaptors. (**B**) Overview of the single cell poly(A) site usage (scPASU) pipeline. Details are described in Materials and Methods. (**C**) Distributions of normalized distances between the transcription start site (TSS) and the mid-point of its poly(A) site(s), where 0.0 is the TSS and 1.0 is the end of the transcription unit. The top graph contains poly(A) sites in human ureter urothelial cells as determined by scPASU; the bottom contains poly(A) sites in the PolyASite atlas (Homo sapiens v2.0, GRCh38.96). (**D**-**E**) UCSC genome browser views for *OCLN* (**D**) and *QSOX1* (**E**) showing the data moving through the scPASU pipeline. From top to bottom: reads after step 1a (Dedup & Uniquely mapped reads), reads after step 1b (Genomic-A filtered reads), reads overlapping with the scPASU poly(A) sites (Filtered reads), urothelial-specific poly(A) sites at the end of step 3 (Poly(A) peaks), signals from PolyASite atlas (PolyASite 2.0), and annotated transcripts from GENCODE V46. See also Figure S1 and Table S1.

Leveraging scPASU, we first established a ureter urothelial cell-specific poly(A) site reference (**Table S1**) and benchmarked it against an existing poly(A) site database, PolyASite 2.0^3^. PolyASite 2.0 contains bulk poly(A)-seq data from various cell lines, blood cell types, and heterogenous tissues but not urothelial cells or their specific differentiation states. Compared with PolyASite 2.0 data, our urothelial cell-specific poly(A) sites demonstrated a comparable distribution across genes (**Fig. 1C**), contained one of 13 known poly(A) signals (**Fig. S1A**), and displayed similarly skewed nucleotide frequencies (**Fig. S1B**). Our poly(A) sites also showed an expected enrichment for 3’ untranslated regions (3’ UTRs) (**Fig. S1C**). Furthermore, we observed remarkable positional overlap between poly(A) sites detected by scPASU and either PolyASite 2.0 data or annotated transcription end sites (TES). For example, poly(A) sites 2 (P2) and 3 (P3) for occludin (*OCLN*) matched the ends of several annotated transcripts (**Fig. 1D**), while P1 for quiescin sulfhydryl oxidase 1 (*QSOX1*) encompassed an annotated TES and P2 matched signals in PolyASite 2.0 (**Fig. 1E**). The same could be illustrated by epidermal growth factor receptor (*EGFR*) (**Fig. S1D**) and *C4orf19* (**Fig. S1E**). Globally, 70.0% of our urothelial-specific poly(A) sites were within 50 nucleotides of a site in PolyASite 2.0, with 69.4% directly overlapping sites in the database (**Fig. S1F**). These observations gave us confidence to proceed with APA testing for differences between specific urothelial cell differentiation states.

## Urothelial cell differentiation-associated APA were distinct from differentially expressed genes and could be spatially resolved *in situ*

Conventional scRNA-seq analysis previously grouped ureter urothelial cells into 8 clusters based on overall gene expression (**Fig. 2A**, top)^1^. These cell clusters could collapse into 3 well-defined differentiation states: basal (clusters 2, 3, and 6), intermediate (clusters 0, 1, and 5), and umbrella (clusters 4 and 7) cells (**Fig. 2A**, bottom). These differentiation states could be readily visualized on a ureter cross section with the basal cell layer abutting the lamina propria and the umbrella cells lining the lumen (**Fig. 2B**). Basal cells were established as the urothelial progenitor cells and were characterized by high keratin 5 (*KRT5*) and exclusive sonic hedgehog (*SHH*) expression^1^. As basal cells differentiate into umbrella cells, the expression of basal cell markers decrease while uroplakin genes, keratin 8 (*KRT8*), and keratin 18 (*KRT18*) expression increase.

**Fig. 2.**
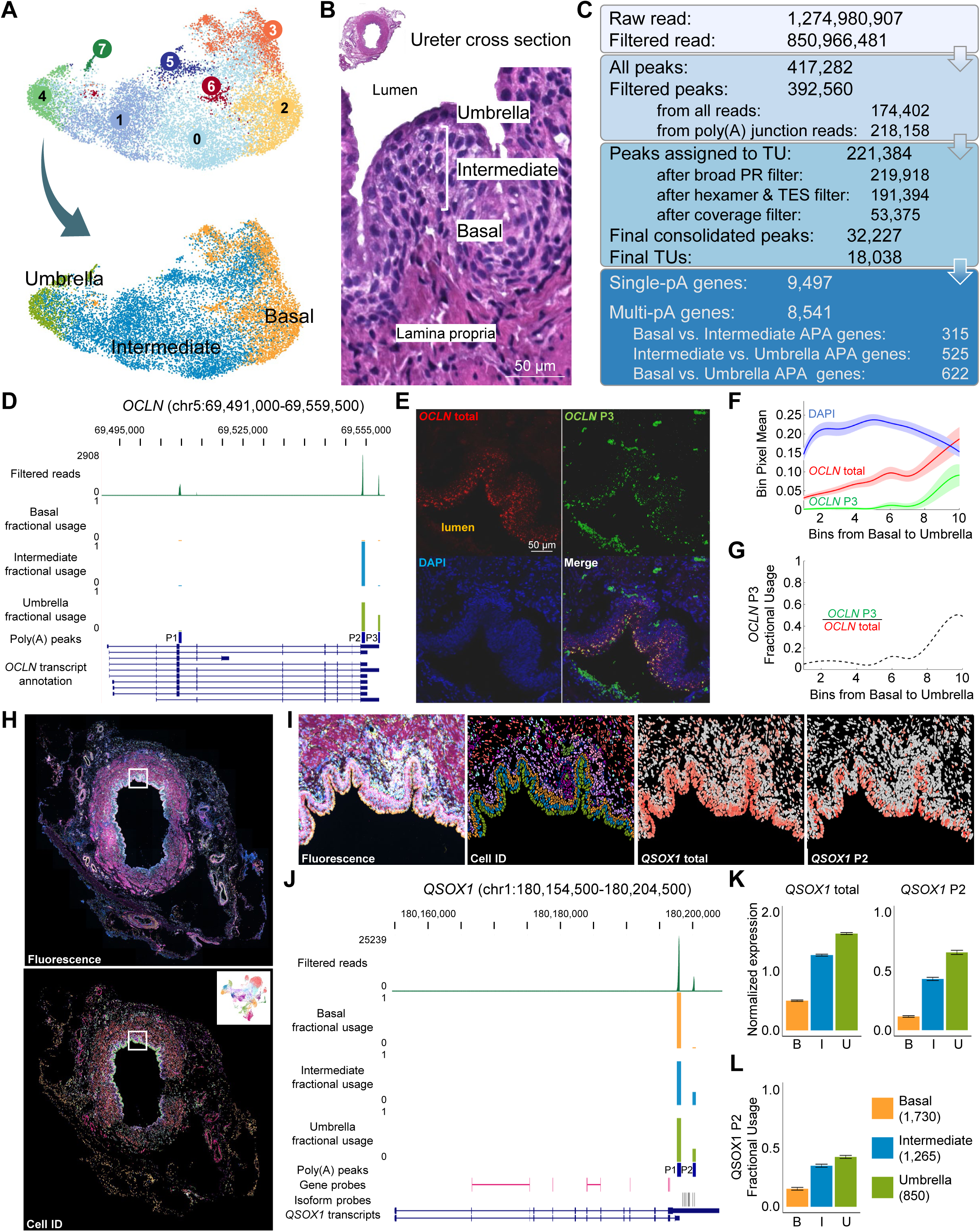
Urothelial cells have differentiation-specific alternative poly(A) site usage. (**A**) UMAP visualization of ureter urothelial cells based on scRNA-seq analysis from Fink and Sona *et. al.* ^1^ (top) and after collapsing cell clusters that made up the basal (clusters 2, 3, and 6), intermediate (clusters 0, 1, and 5), and umbrella (clusters 4 and 7) cell layers (bottom). (**B**) H&E staining of a ureter cross section (top) and a magnified area showing the urothelium with the basal, intermediate, and umbrella cell layers labeled (bottom). (**C**) Summary statistics for each data processing step in the scPASU pipeline for the ureter urothelial cell dataset. (**D-G**) Alternative poly(A) site usage in *OCLN* between intermediate and umbrella cells as detected by scPASU (**D**) and independently validated by RNA FISH immunofluorescence (**E**) and quantification (**F** & **G**). The fractional usage tracks in the UCSC genome browser panel (**D**) show the relative expression of each poly(A) isoform in the specified cell layer. RNA FISH probes were designed against all *OCLN* isoforms (*OCLN* total in red) or specifically for the P3 isoform (*OCLN* P3 in green) in **E**. (**F**) The urothelium was segmented and partitioned into ten concentric bins from the basal cell layer to the umbrella cell layer. The image shows the average pixel intensities per bin for the red (*OCLN* total), green (*OCLN* P3), and blue (DAPI) channels. Average pixel intensities were normalized to a 0-1 scale. Fractional usage of *OCLN* P3 was then reported across the urothelium from basal to umbrella, dividing the cumulative *OCLN* P3 signal intensities by *OCLN* total (**G**). (**H**) Total fluorescence signal from Xenium *in situ* assay on a ureter cross section using a custom probe panel (top), and cell identities from the Xenium data analysis in Fig. S2E (same as the inset UMAP) are projected onto the same cross section (bottom). (**I**) A magnified area (marked by the white rectangle in **H**) of the ureter showing total fluorescence signal, cell identities (basal in orange, intermediate in blue, and umbrella in green), probe signal for all *QSOX1* isoforms (*QSOX1* total), and probe signal for *QSOX1* P2 isoform (*QSOX1* P2). (**J**) UCSC genome browser view for *QSOX1* showing the sequencing reads supporting the poly(A) sites (Filtered reads), fractional usage tracks for each cell layer, location of the poly(A) sites [Poly(A) peaks], and Xenium custom probe positions for total (Gene probes) and P2 isoform (Isoform probes). **K**) Normalized expression of *QSOX1* total and *QSOX1* P2 for the basal, intermediate, and umbrella cells. (**L**) Fractional usage of *QSOX1* P2 for the basal, intermediate, and umbrella cells, computed by dividing *QSOX1* P2 by *QSOX1* total expression for each cell state and calculating the average fractional usage. See also Figure S2 and Table S2.

After enumerating the per cell usage of poly(A) sites in our urothelial cell-specific reference, we performed pair-wise testing for APA in 8,541 genes with multiple poly(A) sites (heretofore referred to as multi-pA genes) among the three differentiation states (**Fig. 2C**). There were 315 APA genes between basal and intermediate cells, 525 between intermediate and umbrella cells, and 622 between basal and umbrella cells (**Fig. S2A** & **Table S2**). Importantly, within each comparison, more than half of the genes showing APA were distinct from those genes that were differentially expressed (**Fig. S2B**). Consequently, these APA genes would have been overlooked by previous analyses as potential biological signals involved in urothelial differentiation.

Next, we sought to validate our identified APA genes using orthogonal assays. OCLN protein is a constituent of tight junctions, which are characteristic of umbrella cells and integral to epithelial barrier function. Concordantly, only intermediate and umbrella cells, but not basal cells (no fractional usage), showed *OCLN* mRNA expression (**Fig. 2D**). Moreover, scPASU identified three poly(A) sites for *OCLN*. Of these, P2 was robustly expressed in both intermediate (fractional usage = 0.997) and umbrella (fractional usage = 0.646) cells while P3 was exclusively detected in umbrella cells (fractional usage = 0.351) (**Fig. 2D** & **Table S2**). We performed RNA-fluorescence *in situ* (RNA FISH) assay on ureter cross sections with probes against all *OCLN* transcripts (*OCLN* total) and probes specific for the P3 isoform (*OCLN* P3) (**Fig. 2E** & **Fig. S2C**). We quantified the fluorescence signals for DAPI, *OCLN* total, and *OCLN* P3 by stratifying the thickness of the urothelium into 10 bins from the basal to umbrella layer (**Fig. S2D**). While *OCLN* total signal was observed in intermediate and umbrella cells (bins 5-10), *OCLN* P3 signal was most evident in the umbrella cells (bins 9-10) (**Fig. 2F**). Likewise, we saw the highest mean fractional usage for *OCLN* P3 in the umbrella cells (bins 9-10) (**Fig. 2G**). Previous molecular studies of the human *OCLN* 3’ UTR did not include the sequence unique to the P3 isoform^4^; however, studies of the rodent *Ocln* 3’ UTR demonstrated that the binding of HuR protein stabilized *Ocln* mRNA and increased translational output^5^. When we analyzed the *OCLN* P3-unique 3’ UTR sequence (chr5:69,554,657-69,558,305) using the RBPDB database^6^, we recovered 81 predicted HuR (gene symbol *ELAVL1*) binding sites, suggesting that the umbrella cell-specific P3 isoform could be important for ensuring robust OCLN protein expression in these terminally differentiated cells.

We also performed additional APA gene validation using the 10x Genomics Xenium In Situ platform that provides spatial imaging in single cells. In addition to canonical cell marker genes, we also designed probe sets recognizing either a specific poly(A) isoform or all isoforms of the same gene. After clustering cells using marker gene expression (**Fig. S2E**), we projected cell identities onto the ureter cross section for visualization (**Fig. 2H**, bottom & **Fig. 2I**, Cell ID panel). The *in silico* analysis identified two poly(A) sites in *QSOX1*, where P1 showed dominant expression in all urothelial cell layers, while intermediate and umbrella cells displayed increased P2 usage (**Fig. 2J**). We designed probes against all transcripts (Gene probes in **Fig. 2J**) as well as P2 isoform-specific probes (Isoform probes in **Fig. 2J**) for *in situ* detection (**Fig. 2I**, *QSOX1* total and *QSOX1* P2 panels). We stratified total *QSOX1* and P2 isoform expression level by cell states (**Fig. 2K**) and computed P2 isoform fractional usage for each cell state (**Fig. 2L**). Similar to the scPASU analysis outputs, intermediate and umbrella cells had comparable P2 fractional usages, which were both higher compared to basal cells.

APA patterns in dystonin (*DST*) and epidermal growth factor receptor (*EGFR*) were also validated using Xenium In Situ (**Fig. S2F**). It is worth noting that *EGFR* P1 was an unannotated poly(A) site in the first intron, located 1,640 bp downstream from the transcription start site (TSS) (**Fig. S1E**). Both scPASU analysis and Xenium data showed basal cells to have the highest fractional usage for *EGFR* P1. Our validation of APA genes by RNA FISH and Xenium demonstrated exquisite spatial resolution of urothelial differentiation-associated APA for the first time.

## Differentiation-associated 3’ UTR-APA genes shared evolutionarily conserved sequences

We next examined if APA genes were enriched for overarching patterns in line with urothelial cell differentiation. Although select genes, like *OCLN*, could be contextualized in urothelial cell differentiation, gene set enrichment analysis on all APA genes yielded only generic terms such as protein binding (GO:MF) and regulation of primary metabolic process (GO:BP). Therefore, we focused on those APA genes that differed only in 3’ UTR lengths (i.e. UTR-APA genes) because prior studies reported differentiation states and proliferation status to be associated with global 3’ UTR shortening or lengthening in a context-dependent manner^7–10^. To evaluate if this was true in the urothelium, we analyzed the UTR-APA genes (n = 297) using the weighted 3’ UTR expression index (WUI), which numerically summarized the relative usage of each poly(A) site in the 3’ UTR for a given gene at each cell state (**Fig. 3A**). For example, a WUI value of 0 indicated that only the most proximal poly(A) site was used whereas a WUI value of 1 indicated that only the most distal poly(A) site was expressed. When the UTR-APA genes were clustered based on their WUI, 164 genes showed 3’ UTR shortening while 133 genes showed 3’ UTR lengthening from basal to umbrella cells (**Table S3**). Therefore, in urothelial cell differentiation, APA genes exhibited both shortening and lengthening, instead of a global, unidirectional isoform shift.

**Fig. 3.**
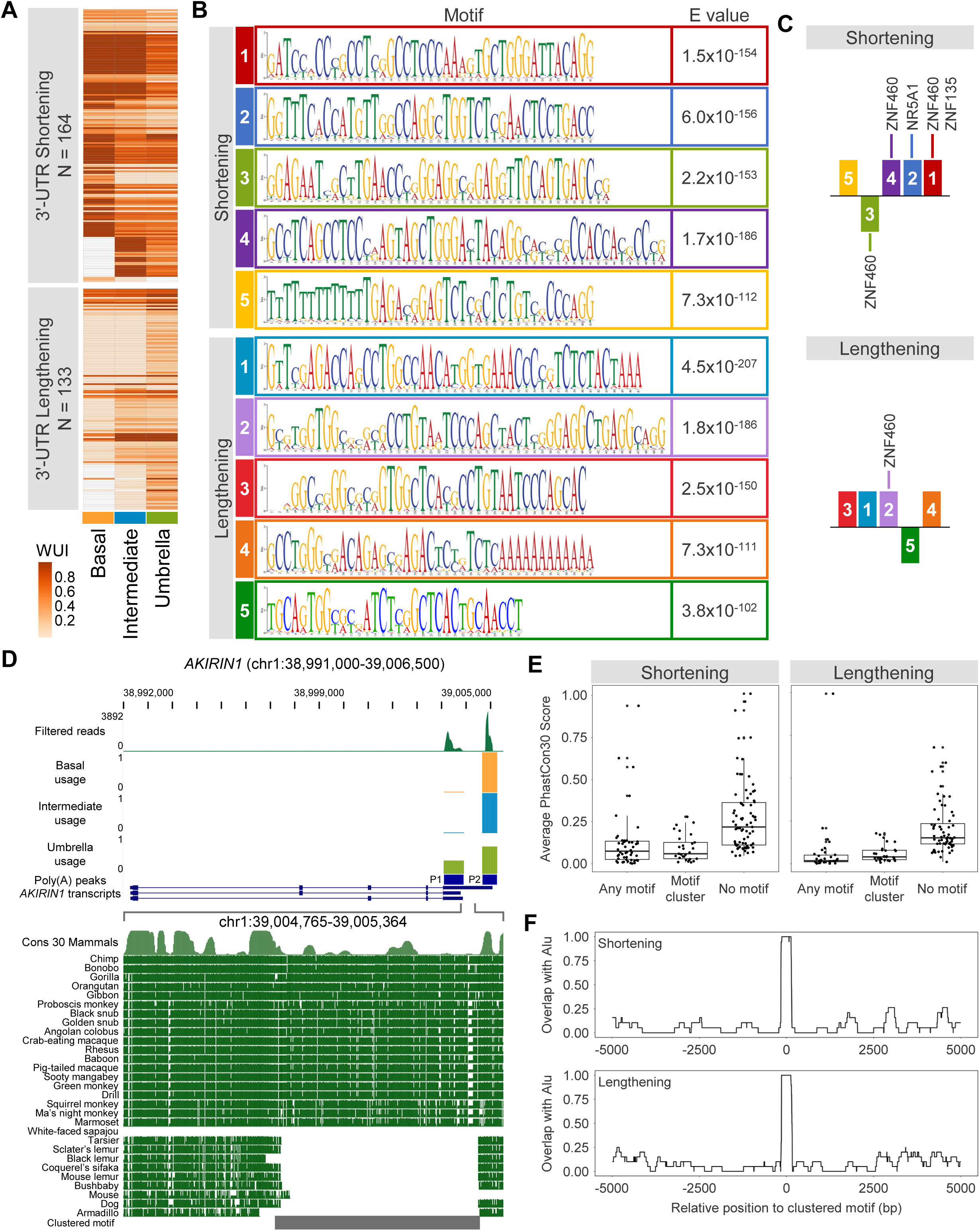
Differentiation-specific 3’ UTR-APA genes in the urothelium have conserved sequence features. (**A**) Heatmaps of weighted 3’ UTR expression indices (WUI) of APA genes with only 3’ UTR poly(A) sites. Genes are grouped into either 3’ UTR shortening (top) or lengthening (bottom) gene sets based on their WUI change along the urothelial cell differentiation axis. (**B**) Top five motifs enriched in the sequence regions between differentially used poly(A) sites of the shortening (top) and lengthening (bottom) gene sets. (**C**) Schematic showing the clustered motifs found in the 3’ UTR sequences of the shortening (top) and lengthening (bottom) gene sets. The motifs are numbered and color-coded as in B. (**D**) UCSC genome browser view for *AKIRIN1*, with a zoomed-in view of the region containing the clustered motif (marked by a gray bar). Poly(A) site reads, cell layer-specific fractional usage tracks, and poly(A) site annotation are shown on the top half. The multiple alignment of 30 mammals against the human reference is shown at the bottom. (**E**) Box plots for average conservation scores for 3’ UTR sequences overlapping with any MEME-discovered motifs (Any motif), those overlapping with full/partial clustered motifs (Motif cluster), and those outside of motifs (No motif) in the shortening (left) and lengthening (right) gene sets. Only the 79 shortening genes and 64 lengthening genes with motifs are plotted. Wilcoxon rank sum test p-values for the shortening gene sets are 3.82 x 10^-8^ (Any motif vs. No motif), 0.851 (Any motif vs. Motif cluster), and 8.08 x 10^-8^ (No motif vs. Motif cluster). Wilcoxon rank sum test p-values for the lengthening gene sets are 4.51 x 10^-11^ (Any motif vs. No motif), 0.005 (Any motif vs. Motif cluster), and 2.97 x 10^-9^ (No motif vs. Motif cluster). (**F**) Distribution of Alu elements in sequences ±5,000 bp from the center of the clustered motif in the shortening (top) and lengthening (bottom) gene sets. Only the 19 shortening genes and 20 lengthening genes whose clustered motif contain an Alu element are plotted. See also Figure S3, Table S3, and Table S4.

Nonetheless, the 3’ UTR is well-recognized for its role in regulating mRNA stability and localization as well as protein translation and trafficking^11–13^. Thus, we assessed the base composition of the 3’ UTR of the UTR-APA genes to identify recurring motifs that might confer a regulatory function. When we analyzed the sequences between the shifting poly(A) sites for possible recurring elements, we saw distinct motif enrichments in the shortening and lengthening gene sets (**Fig. 3B**). We noticed that while the motifs could exist in solitary configurations, they were often observed together in an ordered cluster for both gene sets (**Fig. 3C** & **Table S4**). Intrigued by this unusual arrangement, we concatenated the motifs in the order they appeared in the clusters for the two gene sets separately and then compared the two composite sequences to each other (**Fig. S3A**). Based on sequence alignment, the motif clusters for the shortening and lengthening gene sets had highly similar modular structures and shared substantial sequence identity (149 nucleotides). However, we identified distinct transcription factor binding sites in the divergent sequences of the motif clusters using the JASPAR database (**Fig. 3C** & **Fig. S3A**). Specifically, putative binding sites for zinc finger protein 460 (ZNF460), ZNF135, and nuclear receptor subfamily 5 group A member 1 (NR5A1) in motifs 1, 2, and 3 from the shortening gene set were either completely missing or only partially present in the corresponding motifs of the lengthening gene set. Conversely, in a region of near perfect homology (motif 4 from shortening and motif 2 from lengthening), a significant match with ZNF460 binding sequence was found (**Fig. S3A**). We speculated that transcription factor binding differences might explain why one set of genes shortened their UTRs during urothelial differentiation, but the other set lengthened their UTRs.

While manually examining the UTR-APA genes, we made another unexpected observation about those genes bearing the clustered motifs in their 3’ UTRs. Specifically, the clustered motifs overlapped with sequences that are conserved exclusively in higher primates (**Fig. 3D** & **Fig. S3B**). We quantified this by plotting the average conservation scores across 30 mammals for 3’ UTR sequences overlapping enriched motifs not in clusters (Any motif), those under the motif clusters (Motif cluster), and those outside of any motifs (No motif) (**Fig. 3E**). In this analysis, sequences conserved across the 30 mammals, including mouse, dog, armadillo, and prosimians, would have a high conservation score while sequences conserved in a subset of species, such as in higher primates, would have a low conservation score. Consistent with our manual inspection, sequences in the “Motif cluster” group had significantly lower conservation scores compared to the “No motif” group for both shortening and lengthening gene sets (**Fig. 3E**).

A major demarcating event during primate evolution was the burst of retroelement insertion, dubbed the retrotranspositional explosion, that occurred as prosimians and simians diverged^14^. Given that the motif clusters showed conservation within simians, we looked for similarities between these sequences and retroelements. Focusing on the 29 shortening and 30 lengthening genes with motif clusters in their 3’ UTRs, we assessed the proportional overlap with any repeat sequences for the motif cluster regions and for regions outside of the motif clusters. The motif cluster regions had significantly higher overlap with repeat sequences in general (**Fig. S3C**), and of these, *Alu* elements were found in 19 out of 20 shortening genes and 20 out of 20 lengthening genes (**Fig. S3D-E** & **Table S4**). These *Alu* elements, which spanned the entire motif cluster region in each gene, existed largely as singletons in the 3’ UTRs (**Fig. 3F**). Finally, several of these genes have been shown to promote differentiation in embryonic stem and mesenchymal stem cells as well as other progenitor cells^15–26^. Their discovery here as urothelial differentiation-associated APA genes provides strong impetus to investigate their regulation and function in the urothelium in future studies.

## Discussion

The urothelium of the human ureter represents a highly specialized tissue that must maintain barrier integrity through proper differentiation into a protective umbrella cell layer. Understanding how normal urothelial differentiation is maintained during homeostasis and cycles of injury and repair has a broad impact on advancement in tissue engineering and disease management. A previous scRNA-seq study from the GenitoUrinary Development Molecular Anatomy Project (GUDMAP) consortium revealed that overall gene expression patterns were highly similar among urothelial cell states across differentiation ^1^. Here, we demonstrate that APA contributes to transcriptomic diversity during cellular differentiation and homeostasis. By repurposing 3’ scRNA-seq data and developing a novel computational pipeline, we have uncovered a role for APA in modulating transcript diversity independently of total gene expression, highlighting its role in regulating urothelial differentiation.

First, we established that the urothelial cell-specific poly(A) sites enumerated by our scPASU analysis pipeline showed similar characteristics displayed by *bona fide* poly(A) sites. Our analysis identified differentiation state (basal, intermediate, and umbrella)-specific expression of APA isoforms. Importantly, the majority of such APA genes were distinct from those that were differentially expressed, thus identifying previously unappreciated transcriptome dynamics contributing to cellular differentiation in the urothelium. Furthermore, by leveraging RNA FISH and Xenium In Situ assays, we demonstrated, for the first time, a spatially resolved shift in poly(A) isoform expression. These results point to APA as an additional layer of regulation that ensure cells can produce functionally distinct transcripts without altering overall levels of gene expression.

APA genes are commonly stratified into coding region APA, where the alternative poly(A) isoforms harbor different numbers of exons yielding C-terminally distinct protein isoforms, and UTR-APA, where the alternative isoforms differ only in their 3’ UTR lengths. Both types of APA genes were discovered in our study, but gene set enrichment analysis did not identify coordinated APA regulation of a specific pathway or process. However, many APA genes did have urothelium-specific functions, and their cell state-specific poly(A) isoform expression pattern could explain discrepancies between mRNA and protein levels. For example, OCLN is a tight junction protein expressed mainly in the terminally differentiated umbrella cells to support the barrier function in this luminal cell layer, yet its mRNA is robustly detected in both intermediate and umbrella cells. We found that the umbrella cells uniquely expressed the *OCLN* P3 isoform, which had an extended 3’ UTR containing putative HuR binding sites known to stabilize the mRNA and increase protein translation in rodents^4,5^. Empirical testing is needed to confirm this for *OCLN* and other gene candidates in human urothelial cells.

One of the most intriguing findings in our study is the discovery of conserved regulatory motifs of UTR-APA genes. After classifying UTR-APA genes as shortening or lengthening gene sets based on if a gene had more proximal or distal poly(A) isoforms in umbrella cells compared to basal cells, we found that both shortening and lengthening gene sets showed enrichment of motifs that were conserved only in higher primates. Motifs from shortening and lengthening gene sets differed from each other, which suggests that differential transcription factor binding to the 3’ UTR of these genes may result in differentiation state-specific poly(A) site usage. A significant number of these motif clusters were *Alu* elements, which are primate-specific retrotransposons that appeared between 50 and 25 million years ago when simians and prosimians split^14,27^. Notably, ZNF460 and ZNF135 were among the transcription factors predicted to bind to these sequences, and zinc finger transcription factors were thought to have co-evolved with *Alu* elements^28^. Finally, several APA genes, including *AKIRIN1*, *DNM1L*, and *OPTN*, that contain *Alu* elements in their 3’ UTR have established roles in cellular differentiation outside of the urothelium^15–26^. Altogether, our discoveries here raised an exciting possibility that these 3’ UTR *Alu* elements may have been domesticated to regulate APA and that the targets of such regulation play direct roles in shaping transcriptomic diversity during urothelial differentiation in humans and other higher primates.

Our study established that APA is a mechanism for gene regulation during urothelial differentiation and generated a catalogue of urothelial differentiation-associated APA isoforms. These isoforms can be further studied to better define their function and regulation to advance our understanding of cellular differentiation as well as transcriptome and proteome regulation.

## Methods

### Single cell polyadenylation site usage (scPASU) analysis

#### 1. Data preprocessing (Step 1 in ***Fig. 1B***)

Only sequencing reads belonging to the 13,544 urothelial cells from the original dataset ^1^ were analyzed. After alignment to hg38/GRCh38 using cellranger (v7.1.0), PCR duplicates and ambiguously mapped reads were removed using the *dedup* module of umi_tools (v.1.1.4) ^29^ and samtools ^30^ respectively. Reads likely primed by genomically encoded A-rich sequences were also removed using polyAfilter ^31^. These processed BAM files were converted back to FASTQ files for downstream steps.

#### 2. Peak calling and retention (Step 2 in ***Fig. 1B***)

We used MACS2 ^32^ to identify clustered reads, or peaks, which represented potential poly(A) sites. Peaks with multiple summits were split if gaps in the read alignment were evident between summits. Separately, we identified a small proportion of poly(A) junction reads (164,394,247 out of 1,274,980,907 deduplicated, uniquely mapped reads) to contain the poly(A) junction and partial poly(A) tail using samToPolyA.pl with *minClipped* set to 9 and *minAcontent* set to 0.85 (https://github.com/julienlag/samToPolyA). Only those peaks supported by ≥3 poly(A) junction reads were retained. The poly(A) processing region (PR) for each peak was then inferred from the poly(A) junction reads. In parallel, we also repeated peak calling with just the poly(A) junction reads to rescue any low-coverage poly(A) sites that might have fallen below the noise threshold of MACS2. The two peak sets were then combined to produce a unified peak set for further filtering.

#### 3. Peak clean up and assignment (Step 3 in ***Fig. 1B***)

The unified peak set was further filtered based on gene annotations, inferred PR width, presence of poly(A) signal, and estimated usage. For filtering peaks based on gene annotations, we first created a Transcription Unit (TU) reference by collapsing the transcripts for each gene in the GTF file used for cellranger alignment to create a distinct TU interval for each gene. A 5-kb extension in the 3’ direction (flank region) was also added to each TU to capture potentially unannotated transcription end sites (TES). The final lengths of these flank regions were updated based on the 3’ most peaks for each TU. Peaks were then assigned to either the gene body or the flank region of a TU based on positional overlap (Step 3a). Peaks mapping to more than one TU or no TU and peaks with an inferred PR > 1 kb were removed (Step 3b).

To minimize false positives, we required a peak to either contain at least one poly(A) signal hexamer within or 50 bp upstream from its inferred PR or be within 100 bp from an annotated TES (Step 3c). Lastly, the per gene read coverage was calculated for each peak using the *featureCounts* module of RSubread (v.2.12.3) ^33^, and lowly used peaks (<10%) were filtered out (Step 3d). The final retained peaks were re-assigned to TUs, numbered from 5’ to 3’, and annotated for their genomic contexts (placed in order of priority to resolve ambiguous classifications: 3’ UTR, Transcription Start Site (TSS)-proximal, Exonic, Intronic, and Flank) (Step 3e). We also flagged peaks that were likely to result from exome peaks being split due to spliced alignments. These final annotated peaks represented our dataset-specific poly(A) site reference (**Table S1**).

#### 4. Poly(A) count-by-cell matrix generation (Step 4 in ***Fig. 1B***)

The FASTQ files from step 1 were fed into cellranger (v7.1.0) along with the poly(A) site reference to produce the peak count by cell matrix.

#### 5. Alternative poly(A) site usage testing (Step 5 in ***Fig. 1B***)

A DEXSeq (v.1.44.0) based pipeline, previously developed by our lab for alternative poly(A) (APA) testing on bulk poly(A)-seq data ^2^, was repurposed for APA testing in scPASU. Only TUs with more than one peak (multi-pA genes) were included in APA testing with the criterion that in both testing groups (i.e. the two distinct cell groups), at least one peak must have a read count in at least 10% of the cells. Pairwise testing was performed between basal (clusters 2, 3 and 6), intermediate (clusters 0, 1 and 5), and umbrella cells (clusters 4 and 7). DEXSeq required at least two replicates per group ^34,35^. To accommodate this requirement, three pseudo-replicates were generated by randomly sampling 70% of the cells without replacement. The fractional usage of each poly(A) site in each cell group was calculated by taking the average across its pseudo-replicates. Benjamini-Hochberg correction was applied to the p-values. A peak was considered differentially expressed if it had an adjusted p-value <0.01, an absolute differential fractional usage fold change ≥1.5, a fractional usage per group ≥0.05, and an absolute fractional peak usage difference ≥0.1 (**Table S2**).

##### Characterization of poly(A) sites

The PRs plus 50 bp upstream sequences were scanned for poly(A) signals, and the most downstream occurrence for each poly(A) site was recorded. The same search was performed on a subset of peaks with maximum PR width of 100 bp. This was done to match the size distribution of PRs mapped by bulk poly(A)-seq. For comparison and as a positive control, the same analysis was done for the 50 bp-window upstream of poly(A) sites in the PolyASite atlas (Homo sapiens v2.0, GRCh38.96) ^3^. The poly(A) sites in the PolyASite atlas that overlapped with each other were clustered prior to scanning for poly(A) signals. As a negative control, a null sequence set was created by drawing random sequences from the reference genome matching the size distribution of scPASU inferred poly(A) sites with a maximum PR width of 100 bp. We also plotted the per base nucleotide frequencies for the ±100 bp-region centered on the PR start coordinates in our poly(A) site reference as well as those from the PolyASite atlas. Lastly, a metagene plot showing the distribution of poly(A) sites along the length of their TUs, where 0 denoted the transcription start site and 1 denoted the transcription end site (TES), was created for both our reference and the PolyASite atlas.

##### Differential gene expression testing

We performed pair-wise pseudo-bulk differential gene expression testing between basal, intermediate, and umbrella cells using DESeq2 (v.1.38.3) ^36^. For each comparison, only genes having a read count in at least 10% cells in one cell group were tested. Three pseudo-replicates were generated for each group by randomly sampling 70% of the cells without replacement. We used the Linear Ratio Test of DESeq2, where a Gamma-Poisson Generalized Linear Model ^37^ was fitted and counts were normalized using the standard median ratio method introduced in DESeq ^38^. Benjamini-Hochberg correction was applied to the p-values. A gene was considered differentially expressed if it had an adjusted p-value <0.01 and an absolute fold change ≥1.5.

##### Xenium spatial gene expression assay and data analysis

###### Gene panel design

A custom Xenium human lower urinary tract probe panel targeting 341 genes and mRNA isoforms, including those related to urothelial, fibroblast, immune, and smooth muscle cell markers and negative controls for cell type identification, was designed under panel ID #8QH2R6.

###### Xenium sample preparation

A 10 µm, OCT-embedded frozen ureter sample from a deceased organ donor was sectioned onto a Xenium slide and stored at -80°C. After thawing the slides at 37°C for 1 min, it was fixed in 4% PFA for 30 min and washed in PBS. The Slide was then permeabilized using 1% SDS for 2 min, followed by 2 PBS washes and immersed in pre-chilled 70% methanol for 60 min on ice and washed 2 times in PBS. A custom probe panel and control probes were added to the slide at a concentration of 10 nM and hybridized at 50°C overnight in a thermocycler. Stringent post-hybridization washes were performed, followed by ligation of probes at 37°C for 2 hr and primer annealing. Rolling circle amplification of probes was performed for 2 hr at 37°C, thereby generating copies of the gene-specific barcode for each RNA-binding event. After washing and chemically quenching the background fluorescence, nuclei were stained with DAPI, and the slide was loaded onto a Xenium Analyzer.

###### Xenium Analyzer

Tissue sections were incubated with reagents and fluorescent-labeled probes for detecting RNA provided by manufacturer. The instrument automatically cycled in reagents, imaged the sections, and removed reagents for 15 rounds. Across the entire tissue thickness, Z-stack images spanning a 0.75-μm step size were acquired.

###### Image processing and analysis

A single image representative of 1 region of interest (ROI) was created using Z-stack images taken in every cycle and channel and stitching on a DAPI image. This image was used to build a spatial transcriptomic map of the tissue section. Xenium Onboard Analysis pipeline detected all potential mRNA as ‘puncta’, which is defined as observed photons in every cycle, image and color. The observed emitted light across different cycles from localized puncta were fitted into a Gaussian distribution and a unique optical gene signature was generated after 15 Xenium cycles. For each decoded transcript, a Q score was assigned and transcripts with Q ≥ 20 were considered for Xenium spatial gene plots. The cell segmentation was calculated in a 15-μm radius from the nucleus outward or until a boundary was reached based on DAPI staining to generate a feature-cell matrix.

###### Post-Xenium histology

H&E staining was performed on the same tissue sections that underwent Xenium analysis. A Keyence (Itasca, IL) microscope was used to image the slide at 20x magnification, and images were stitched together to generate a whole-slide bright-field image.

###### Data QC

The raw Xenium ureter data, which included 78,683 cells and 8,916,871 total transcripts (94 median transcripts/cell, 29 median genes/cell) was filtered for low-quality cells using QC thresholds on the number of genes (≥5) and transcripts (≥80) per cell. The final data included 43,683 quality cells, with medians of 149 transcripts per cell and 40 genes per cell, for downstream analysis.

###### Cell clustering and identification

Downstream analysis was completed in R (v4.3.0) using the Seurat (v5.1.0) standard workflow for analysis of image-based spatial data. The Xenium assay was normalized, principal components (PCs) were computed using genes only (isoforms excluded), and clustering optimization was completed using clustree ^39^ to fine tune the number of PCs and resolution. Isoforms were excluded from clustering optimization so as not to bias urothelial clustering and downstream fractional usage calculations. Using 30 PCs and 0.4 resolution, 18 clusters were visualized using UMAP and annotated as belonging to one of three compartments –urothelium, stromal, or immune– based on marker gene expression. Marker genes included KRT5 and UPK3A for urothelial cells, ACTA2, PDGFRA, PECAM1, XKR4, and RGS5 for stromal cells, and PTPRC for immune cells. Subset analysis was then performed for each of the three compartments following the same Seurat workflow.

For each subset analysis, cells from clusters assigned to the urothelial, stromal, or immune compartments were processed by the pipeline as described above, starting from normalization and clustering optimization. Optimal clustering parameters for each compartment included 5 PCs and 0.1 resolution for urothelial cells, 13 PCs and 0.3 resolution for stromal cells, and 7 PCs and 0.2 resolution for immune cells. As a result, 3 urothelial clusters (3,845 total cells), 11 stromal clusters (33,702 total cells), and 4 immune clusters (6,136 total cells) were identified. To determine the identity of each compartment’s clusters, marker gene expression was visualized as feature plots.

###### Isoform fractional usage

We also computed the fractional usage of several significant APA isoforms for *QSOX1*, *DST*, and *EGFR*. For *QSOX* P2 and *DST* P2, per cell fractional usage was computed as the normalized expression of isoform divided by normalized expression of total gene per cell. For *EGFR* P1 fractional usage was computed per cell as the normalized expression of isoform divided by the sum of normalized expression of isoform plus normalized expression of total gene. The average across all cells was then computed to provide the overall fractional usage for a given cell population.

##### RNA FISH assay and quantification

In situ hybridization was performed using RNAscope following the manufacturer’s protocol (ACDbio) using their supplies unless otherwise indicated. Briefly, human ureters were fixed overnight in 4% paraformaldehyde at 4°C, embedded in paraplast, and sectioned at 10 µm. FPE sections were deparaffinized in xylene, followed by a dehydration into distilled water using an ethanol series. Sections were then incubated in RNAscope-H_2_O_2_ for 10 min at room temperature followed by 15 min at 98°C in RNAscope-Target-Retrieval reagent and 30 min at 40°C in Protease I solution. Sections were then hybridized with the RNAscope buffered Z probes for *OCLN* (ACDbio ref. 406511, detecting the open reading frame of *OCLN*), and *OCLN-O2-C2* (ACDbio ref. 1289781-C2, detecting the 3’ UTR region specific for *OCLN* P3 transcript) for 2 h at 40°C and stored overnight in 5x SSC. The next day, sections were sequentially incubated in AMP1 for 30 min at 40°C, AMP2 for 30 min at 40°C, and then AMP3 for 15 min at 40°C to amplify the probes. To visualize the individual channels, the sections were incubated with a channel-specific HRP and then with either the red dye TSA^+^Cy3 (Akoya Biosciences #NEL744001KT) or the far-red dye TSA^+^Cy5 (Akoya Biosciences #NEL745001KT) for 30 min at 40°C. After the incubation with the fluorescent dye, sections were treated with the HRP blocker for 20 min at 40°C before continuing to the next channel. After a final wash, sections were incubated with DAPI and embedded in ProLong™ Diamond Antifade Mounting media (Thermo Fisher Scientific). Slides were imaged with a Leica DM5500 using the LASX software and image deconvolution.

Red-Green-Blue (RGB) images of the RNA FISH labeled urothelium were analyzed using computational image analysis in MATLAB. First, the urothelium was segmented manually from the image. The urothelium was then automatically partitioned into ten bins, concentric pixel layers of equal breadth, from basal to umbrella, and the average pixel intensities of each bin for the *OCLN* total (R), *OCLN* P3 (G), and DAPI (B) were computed. Average intensities were normalized between zero and one and plotted as line graphs for each RGB channel. This analysis effectively quantified an intensity profile for each column of the image wherein the urothelium was present (*n* = 670). The fractional usage of *OCLN* P3 was then computed by dividing the cumulative *OCLN* P3 signal intensity per bin by the cumulative *OCLN* total signal intensity.

##### Weighted UTR expression index (WUI) calculation

To classify APA genes undergoing 3’ UTR shortening or lengthening from basal to umbrella cells, we calculated weighted 3’ UTR expression index (WUI) ^40^ for eligible genes, which were multi-pA genes with all poly(A) sites in their 3’ UTR and/or Flank regions. The WUI ranged from 0 to 1, where 0 indicated exclusive usage of the most proximal poly(A) site and 1 indicated exclusive usage of the most distal poly(A) site. After computing the WUIs for genes independently in each cell group, we stratified the genes by their WUI changes across the differentiation spectrum. Genes with decreasing WUIs (n = 164) from basal to umbrella cells were classified as 3’ UTR shortening while genes with increasing WUIs (n = 133) from basal to umbrella were classified as 3’ UTR lengthening (**Table S3**). Each category of genes was then clustered using Pearson correlation to generate the heatmaps.

##### 3’ UTR motif discovery and analysis

For the 3’ UTR shortening and lengthening gene sets, we extracted the sequences between the poly(A) sites undergoing APA from each gene and performed motif discovery using MEME in the “Discriminative mode” to search for the top five motifs with “Zero or One Occurrence per Sequence (zoops)” distribution ^41^. To run the “Discriminative mode”, we generated a length-matched control sequence set that was 10 times larger in numbers (e.g. 1,640 controls sequences for the 164 shortening genes) by sampling from the 3’ UTRs of genes from the same chromosomes as the input gene set. We then compared each of the discovered motif to the JASPAR (Non-redundant) DNA database (JASPAR CORE (2022) vertebrates) using Tomtom ^42^ and reported motif matches with a q-value <0.05 (**Fig. S3A**). The clustered motifs were concatenated separately for shortening and lengthening gene sets and aligned to each other using clustalo (1.2.4) for comparison ^43^.

For conservation analysis, we extracted the phastCons score from Con 30 Mammals by PhastCons ^44,45^ for each 10 bp-bin in the 3’ UTR sequences. We then calculated and plotted the average phastCons scores for a gene across 1) bins overlapping with any MEME-discovered motifs (Any motif in **Fig. 3E**), 2) bins overlapping with full/partial clustered motifs (Motif cluster in **Fig. 3E**), and 3) all bins not overlapping with motifs (No motif in **Fig. 3E**). We considered a gene to contain a partial clustered motif if it had at least 3 out of 5 motifs grouped together in the same order and relative strand orientation as the full clustered motif (**Table S4**). Focusing on the genes containing full/partial clustered motifs, we computed the proportional overlap with annotated repeats (“rmsk.txt” from UCSC genome browser) for 1) sequences overlapping with motifs (Motif cluster in **Fig. S3C**) and 2) sequences outside of motifs (No motif in **Fig. S3C**). Wilcoxon rank sum test was used to determine statistical significance in both the conservation and repeat element analyses.

## Acknowledgements

National Institute of Diabetes Digestive and Kidney Diseases grant U01DK131383 (OW, BHL, AHT)

National Cancer Institute grant R01CA230033 (AHT)

## Author contributions

Conceptualization: SS, AHT

Methodology: NBL, SS, BS, SHL

Investigation: NBL, SS, BS, RAS, VK, WIP, JMY

Visualization: NBL, BS, AHT

Funding acquisition: OW, BHL, AHT

Project administration: AHT

Supervision: OW, KR, AHT

Writing – original draft: BHL, AHT

Writing – review & editing: NBL, SS, BS, VK, KR, SHL, JMY, OW

## Data availability

Ureter scRNA-seq data is available at GEO under accessing number GSE184111. The gene and peak count matrices are available under GSE262863. Codes are available at https://github.com/ninhleba/scPASU_pipeline.

---

**Fig. S1.**
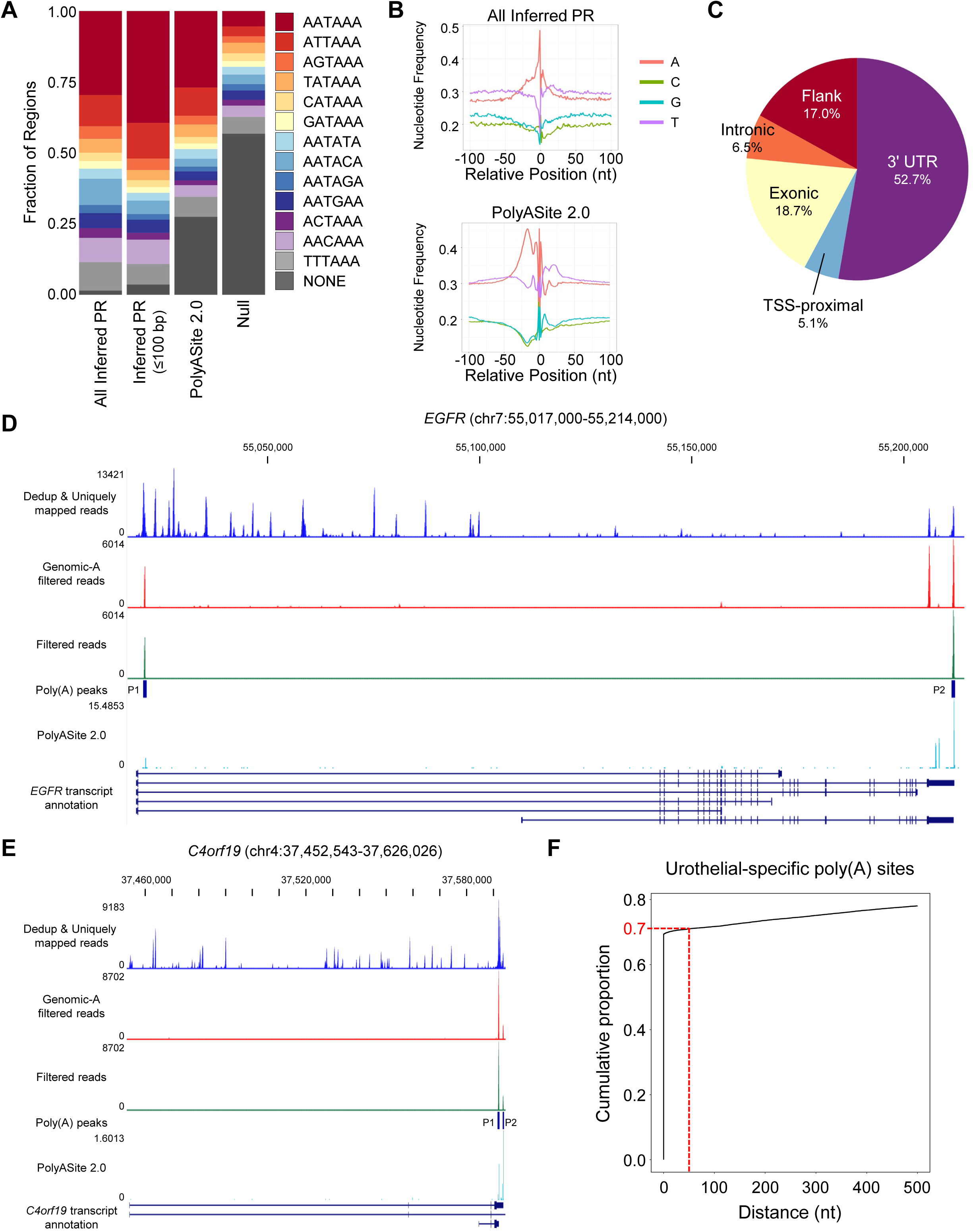
Urothelial-specific poly(A) sites from scPASU have characteristics comparable to known poly(A) sites. (**A**) Frequency plot for poly(A) signal occurrences in the poly(A) processing (PR) regions for all poly(A) sites in the urothelial-specific poly(A) reference (All Inferred PR), the subset of poly(A) sites with inferred PR ≤100 bp [Inferred PR (≤100 bp)], the poly(A) sites in the PolyASite atlas (PolyASite 2.0), and a size-matched random sequence set (Null). (**B**) Nucleotide frequency plot for sequences ±100 bp from the start coordinates of inferred PRs for all poly(A) sites in the urothelial-specific poly(A) reference (All Inferred PR, left panel) and for the PolyAsite atlas (PolyASite 2.0, right panel). (**C**) Genomic contexts for all poly(A) sites in the scPASU-derived urothelial-specific poly(A) reference. (**D**-**E**) UCSC genome browser views for *EGFR* (**D**) and *C4orf19* (**E**) showing the data moving through the scPASU pipeline. From top to bottom: reads after step 1a (Dedup & Uniquely mapped reads), reads after step 1b (Genomic-A filtered reads), reads overlapping with the scPASU poly(A) sites (Filtered reads), urothelial-specific poly(A) sites at the end of step 3 (Poly(A) peaks), signals from PolyASite atlas (PolyASite 2.0), and annotated transcripts from GENCODE V46. (**F**) The cumulative distribution curve of the distance between our urothelial-specific poly(A) sites (n=31,385, only sites associated with TUs that are also represented in PolyASite 2.0 are plotted) and poly(A) sites in PolyASite 2.0.

**Fig. S2.**
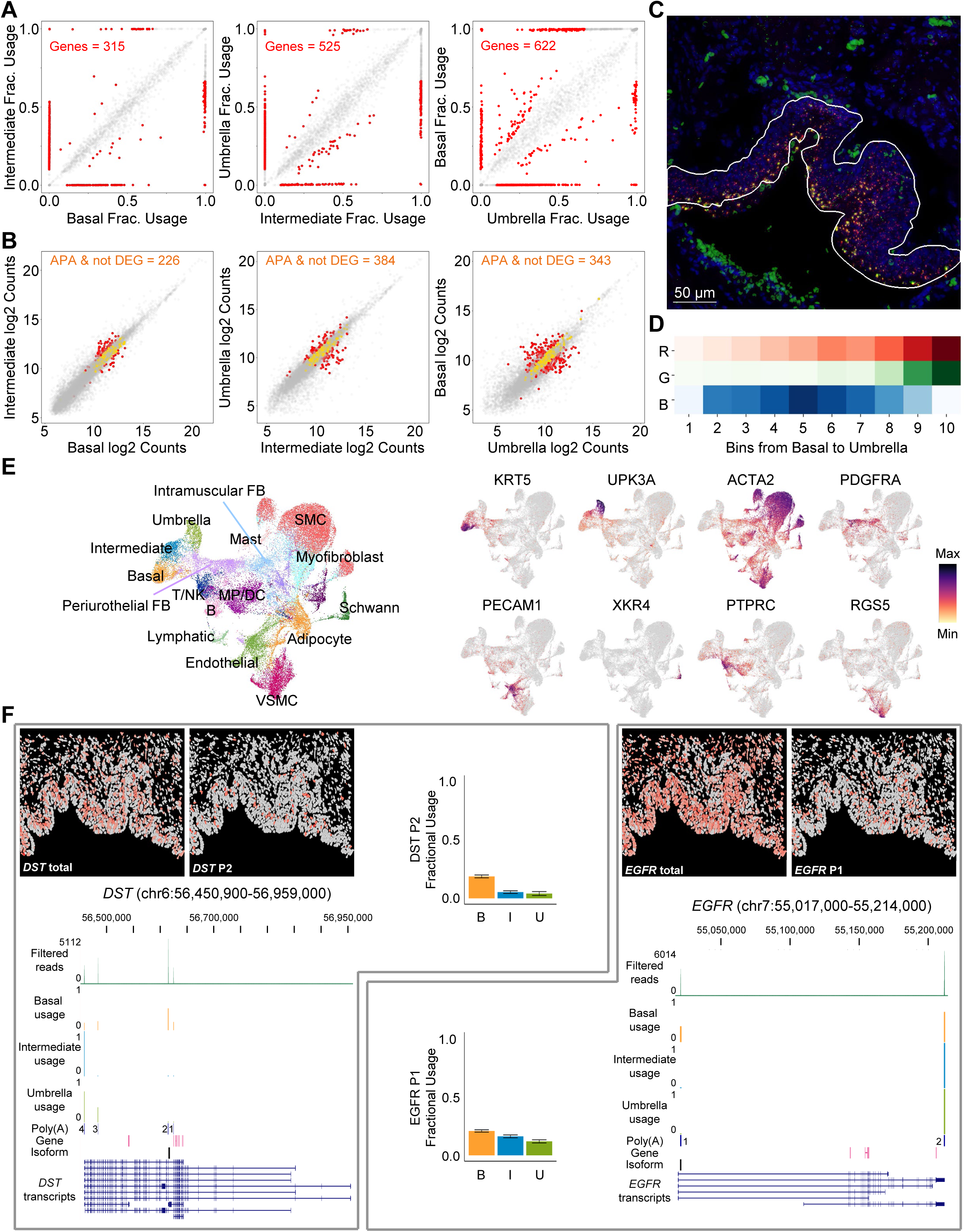
Urothelial differentiation-specific APA events determined by scPASU can be independently validated *in situ*. (**A**) Scatter plots of poly(A) site fractional usages between intermediate and basal cells (left), umbrella and intermediate cells (middle), and basal and umbrella cells (right). Statistically significant APA sites in each comparison are colored in red, with the corresponding gene numbers labeled at the top. (**B**) Scatter plots showing total gene expression comparisons between intermediate and basal cells (left), umbrella and intermediate cells (middle), and basal and umbrella cells (right). Red dots are APA genes that also showed overall differential expression, and orange dots are APA genes that were not differentially expressed between the comparison groups. (**C**) Segmented urothelium used in quantification of RNA FISH *OCLN*, *OCLN* P3, and DAPI intensities. The white outline marks the areas included in the analysis. (**D**) Average bin intensities of *OCLN*, *OCLN* P3, and DAPI computed across the urothelium from basal to umbrella. (**E**) UMAP of the 18 ureter cell populations identified in Xenium data analysis and feature plots for canonical markers used in cell type identification: *KRT5* for basal cells, *UPK3A* for umbrella cells, *ACTA2* for smooth muscle cells, *PDGFRA* for fibroblasts, *PECAM1* for endothelial cells, *XKR4* for Schwann cells, *PTPRC* for immune cells, and *RGS5* for vascular smooth muscle cells. (**F**) Image dimplots show the total gene and isoform expression in each cell in red. Bar plots show the fractional usage of *DST* P2 and *EGFR* P1 for basal, intermediate, and umbrella cells from Xenium data, while UCSC Genome Browser tracks with annotations display the genomic coordinates and fractional usage of each isoform computed by scPASU.

**Fig. S3.**
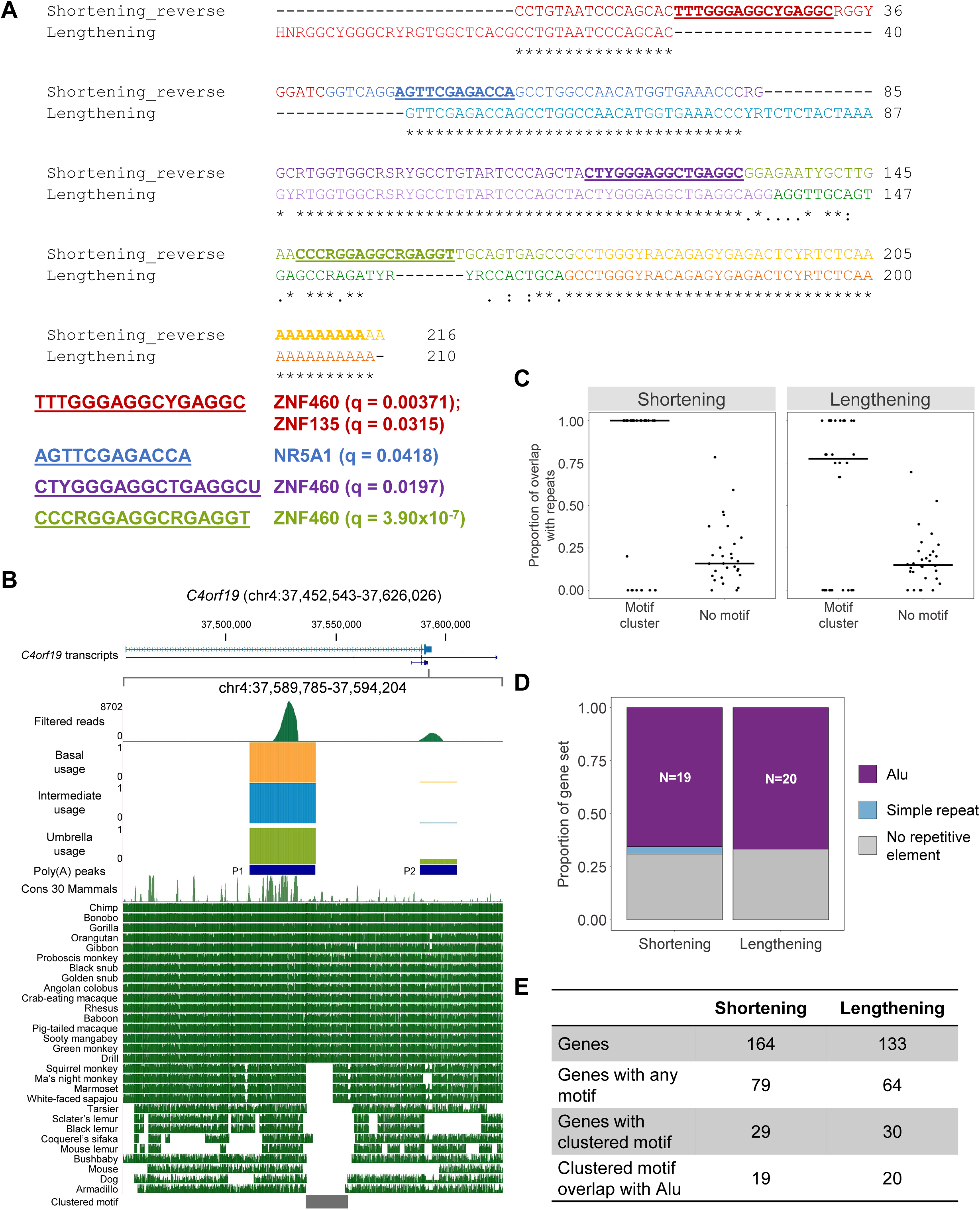
The clustered motifs in 3’ UTR APA genes suggest unique regulation. (**A**) Sequence alignment between concatenated clustered motifs from shortening and lengthening gene sets. Individual motifs are color-coded as in Fig. 3B. The bold underlined bases are sequences that match a transcription factor binding motif in JASPAR CORE (2022) vertebrates database. (**B**) UCSC genome browser view for *C4orf19*, with a zoomed-in view of the region containing the clustered motif (marked by a gray bar). Poly(A) site reads, cell layer-specific fractional usage tracks, and poly(A) site annotation are shown on the top half. The multiple alignment of 30 mammals against the human reference is shown at the bottom. (**C**) Distributions of proportional overlap with annotated repeats for sequences overlapping with full/partial clustered motifs (Motif cluster) and those outside of motifs (No motif) in the shortening (left) and lengthening (right) gene sets. Only the 29 shortening genes and 30 lengthening genes containing clustered motifs in their 3’ UTRs are plotted. Wilcoxon rank sum test p-values between the two groups are 0.0132 for the shortening set and 0.0099 for the lengthening set. (**D**) Bar plots showing the dominant repetitive element that overlapped with each clustered motif in the shortening and lengthening gene sets. (**E**) Summary of the shortening and lengthening gene sets.

